# Age differences in aperiodic neural activity measured with resting EEG

**DOI:** 10.1101/2021.08.31.458328

**Authors:** Ashley Merkin, Sabrina Sghirripa, Lynton Graetz, Ashleigh E. Smith, Brenton Hordacre, Richard Harris, Julia Pitcher, John Semmler, Nigel C. Rogasch, Mitchell Goldsworthy

## Abstract

Previous research using electroencephalography (EEG) and magnetoencephalography (MEG) has shown that neural oscillatory activity within the alpha band (8-12 Hz) becomes slower and lower in amplitude with advanced age. However, most studies have focused on quantifying age-related differences in periodic oscillatory activity with little consideration of the influence of aperiodic activity on these measures. The aim of this study was to investigate age differences in aperiodic activity inherent in the resting EEG signal. We assessed aperiodic activity in 85 healthy younger adults (mean age: 22.2 years, SD: 3.9, age range: 18–35, 37 male) and 92 healthy older adults (mean age: 66.1 years, SD: 8.2, age range 50–86, 53 male) by fitting the 1/f-like background activity evident in EEG power spectra using the fitting oscillations & one over f (FOOOF) toolbox. Across the scalp, the aperiodic exponent and offset were smaller in older compared to younger participants, reflecting a flatter 1/f-like slope and a downward broadband shift in the power spectra with age. Before correcting for aperiodic activity, older adults showed slower peak alpha frequency and reduced peak alpha power relative to younger adults. After correcting for aperiodic activity, peak alpha frequency remained slower in older adults; however, peak alpha power no longer differed statistically between age groups. The large sample size utilized in this study, as well as the depth of analysis, provides further evidence that the aperiodic component of the resting EEG signal is altered with aging and should be considered when investigating neural oscillatory activity.

## 1. Introduction

Aging is associated with various physical, biological, and cognitive changes that can greatly impact one’s physical and mental health. From mid to late life, the brain undergoes age-related changes in both morphology and physiological dynamics, each of which has been linked to changes in behavior and cognition (Harada et al., 2013; Phillips and Andres, 2010). A considerable body of research using electroencephalography (EEG) and magnetoencephalography (MEG) has shown changes in neural oscillatory dynamics with increasing age, especially in the alpha band (∼8–12 Hz) (Babiloni et al., 2006; Grandy et al., 2013; Michels et al., 2013; Scally et al., 2018)t al., 2018; Schaworonkow and Voytek, 2021; Vaden et al., 2012). Individual peak alpha frequency has been found to represent an individual, physiological characteristic that increases from childhood to adulthood, but begins slowing around mid-life (Michels et al., 2013; Scally et al., 2018). Older adults exhibit significantly lower peak alpha parameters (both frequency and power) relative to younger adults (Babiloni et al., 2006; Klimesch, 1999; Sghirripa et al., 2020), with slowing of alpha oscillatory activity and reduced alpha power also observed in multiple age-related dementias, including Alzheimer’s Disease (Neto et al., 2015; Neto et al., 2016), Parkinson’s Disease (Jeong et al., 2016), Lewy Body Dementia (Colloby et al., 2016; Garn et al., 2017), and frontotemporal dementia (Nishida et al., 2011).

In addition to periodic oscillations, recent research has shed light on other components of the EEG broadband signal that could have implications for the interpretation of past findings. Neural oscillations are reflected as “periodic” fluctuations in the EEG signal and are commonly quantified by averaging across fixed frequency bands in EEG power spectra. However, this approach ignores the contribution of “aperiodic” activity to the power spectra, commonly referred to as background noise, which is characterized by a 1/f-like distribution, and changes with task demand and behavioral state (Donoghue et al., 2020; Grigolini et al., 2009; Kumar and Parmananda, 2018; Voytek et al., 2015). Aperiodic activity can be characterized by two parameters: an offset parameter reflecting the uniform shift of power across frequencies and a slope parameter, which delineates the steepness of the 1/f-like function. This slope parameter, often referred to as ‘aperiodic exponent’, is thought to reflect a shift in the excitation/inhibition balance and a decoupling of neuronal spiking activity from the oscillation frequency (Donoghue et al., 2020; Gao et al., 2017; Manning et al., 2009; Miller et al., 2012; Voytek and Knight, 2015; Winawer et al., 2013).

A number of recent studies have demonstrated age differences in aperiodic activity. For example, task-related EEG studies have suggested that as age increases, the aperiodic slope flattens (Dave et al., 2018; Voytek et al., 2015; Waschke et al., 2017), possibly reflecting age-related changes in excitation/inhibition balance (Gao et al., 2017) and an increase in asynchronous neuronal population spiking that is detrimental to cognitive performance (Voytek et al., 2015; Voytek and Knight, 2015). However, it remains largely unknown how these differences in the aperiodic component, if left unaccounted for, might affect the measurement of alpha oscillatory dynamics in aging. A recent study by Donoghue et al. (2020) assessed age differences in resting alpha oscillations, as measured by EEG, after isolating the oscillatory peak from the aperiodic background. Similar to previous studies measuring EEG during task performance, they observed a flatter aperiodic slope and reduced offset in older, relative to younger adults. Peak alpha characteristics also differed between age groups after adjusting for aperiodic activity, with older adults having decreased peak alpha power and a slower peak alpha frequency, although the age difference in aperiodic-adjusted peak alpha power was reduced compared to non-adjusted power.

It is worth noting that the intention of the Donoghue et al. (2020) study was to introduce an algorithm for parameterizing neural power spectra into periodic and aperiodic components, and that analysis of age-related differences in resting EEG spectral parameters were included as one example to demonstrate the algorithm’s utility. Accordingly, the scope of analysis was understandably limited to include just a single channel and a relatively small sample size (16 younger, 14 older). Therefore, the aim of the present study was to extend these findings to consider whole-scalp resting EEG recordings from a much larger sample of younger and older adults. We hypothesized that, in accordance with previous research, aperiodic exponent and offset would be reduced in older adults, and that these age differences in aperiodic activity drive some of the differences in peak alpha characteristics that are otherwise attributed to oscillatory changes.

## 2. Methods

### 2.1 Participants

Data were combined from six similarly designed studies that all included an eyes-closed resting EEG recording, resulting in 85 younger adults (mean age: 22.2 years, SD: 3.9, age range: 18–35, 37 male) and 92 older adults (mean age: 66.1 years, SD: 8.2, age range 50–86, 53 male) in the final sample. All older adults were without cognitive impairment, as assessed using either Mini-Mental State Examination (score >24) (Folstein et al., 1975) or Addenbrooke’s Cognitive Examination (ACE-III) (score >82) (Mioshi et al., 2006). Exclusion criteria for both age groups were history of psychological or neurological disease, a history of substance abuse, medications that alter the function of the nervous system, and an uncorrected hearing or visual impairment. All participants gave written consent and the studies were approved by the University of Adelaide Human Research Ethics Committee, The Queen Elizabeth Hospital Human Research Ethics Committee, and the University of South Australia Human Research Ethics Committee.

### 2.2 EEG data acquisition

Participants were seated in a comfortable chair in a quiet room. They were asked to keep their eyes closed during recording, to remain as still, quiet, and relaxed as possible, and to refrain from actively engaging in any cognitive or mental tasks. EEG data were recorded for 3 minutes from either 57 or 62 electrodes arranged in a 10-10 layout (Waveguard, ANT Neuro, Enschede, The Netherlands) using a Polybench TMSi EEG system (Twente Medical Systems International B.V, Oldenzaal, The Netherlands) or an ASA-lab EEG system (ANT Neuro, Enschede, Netherlands). Conductive gel was inserted into each electrode using a blunt-needle syringe in order to reduce impedance to <5 kΩ. The ground electrode was located at AFz. Signals were sampled at 2048 Hz, amplified 20x, online filtered (DC–553 Hz), and referenced to the average of all electrodes.

### 2.3 EEG pre-processing

Resting EEG data were pre-processed using EEGLAB (Delorme and Makeig, 2004), TESA (Rogasch et al., 2017), and customized scripts in MATLAB (R2019a, The Mathworks, USA). Poor or disconnected channels were removed. Data were then band-pass (1–100 Hz) and band-stop (48–52 Hz) filtered (zero-phase Butterworth, fourth-order) and epoched in 2-second segments. Due to differences in pre-processing pipelines between studies, a subset of data had been down-sampled to 256 Hz prior to epoching (25 younger and 32 older participants; chi-square test, χ^2^_1_ = 0.58, *p* = .44). Independent component analysis was performed using the FastICA algorithm (Hyvärinen and Oja, 2000) and artefacts reflecting eye blinks or scalp muscle activity were identified and removed from the mixing matrix before reconstructing the data. Epochs were then visually inspected for any remaining non-stereotypic artefacts and excluded if necessary. Removed channels were replaced using spherical interpolation.

### 2.4 Spectral analysis

Frontal channels FP1, FPz, FP2, AF7, and AF8 were excluded from datasets collected using 62 electrodes, leaving the same common subset of 57 channels for spectral analysis. Power spectral densities were calculated for each channel using Welch’s method (2-s Hamming windows, 50% overlap). We used the fitting oscillations & one over f (FOOOF) algorithm (Donoghue et al., 2020) to parameterize power spectra into aperiodic and periodic components, extracting the offset and exponent parameters for the aperiodic component across the frequency range 2–40 Hz (knee parameter fixed to 0). Peak alpha frequency (defined as the frequency with largest logarithmic power between 6–13 Hz, detected using MATLAB’s *findpeaks* function) and peak alpha power (the logarithmic power at the peak) were assessed from power spectra at each channel, first without controlling for the aperiodic component (i.e., *‘uncorrected’*), then after subtracting the aperiodic component from the original spectra (i.e., *‘1/f-corrected’*) (Mahjoory et al., 2020) (see Figure 1). If a peak was not observed for a given channel (<0.1% of cases for all participants and channels), then peak alpha frequency and power for that channel were spherically interpolated using EEGLAB’s *pop_interp* function.

**Figure 1.**
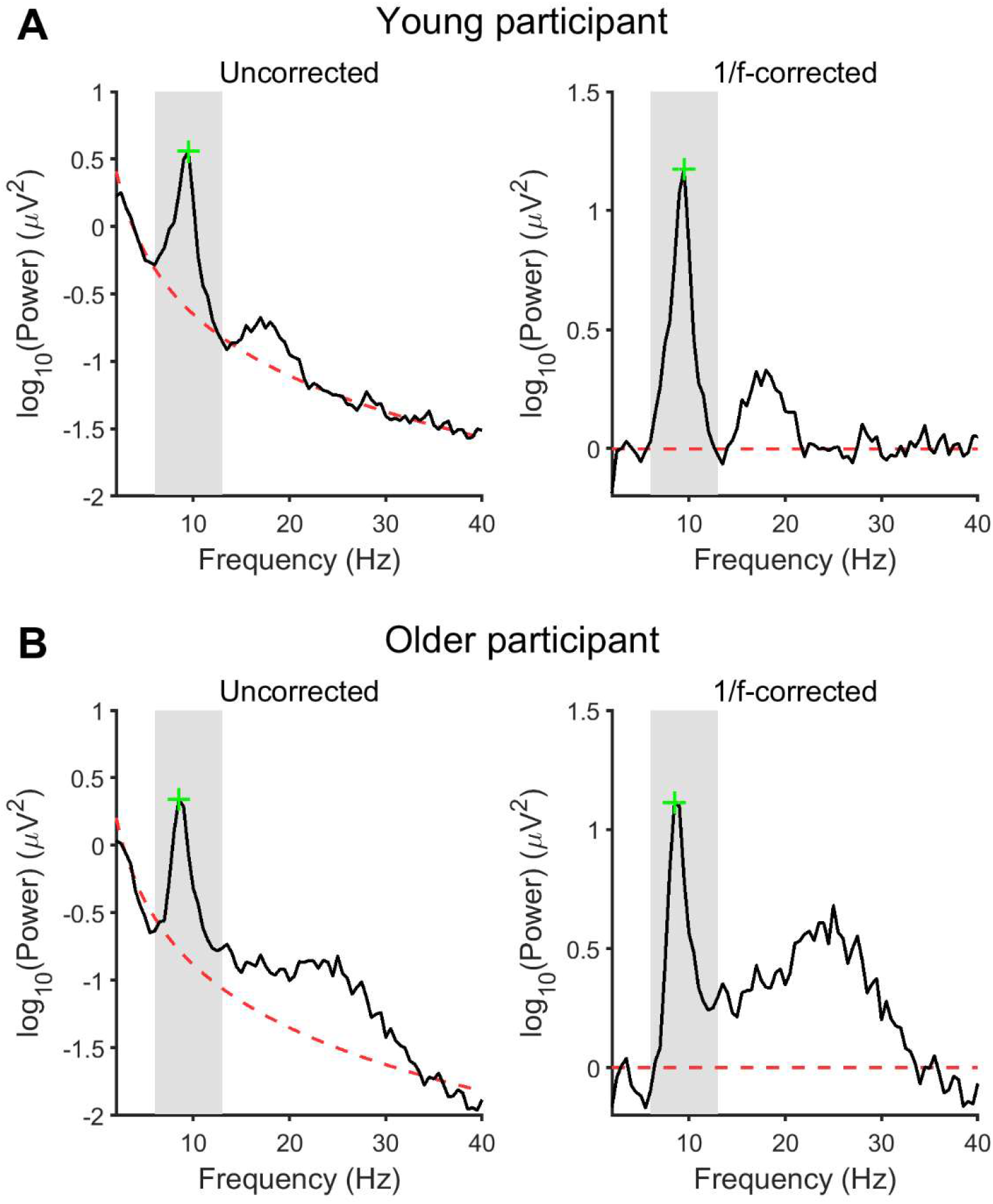
Evaluating peak alpha frequency and power before and after correcting for aperiodic neural activity in the resting EEG power spectrum. Data are from channel Cz in one young (A) and one older (B) participant. The 1/f-like aperiodic component (dashed red line) was estimated from the original power spectrum (left), then subtracted to provide a 1/f-corrected spectrum (right). The grey-shaded region denotes the 6–13 Hz extended alpha range. The green cross denotes the alpha peak.

In addition to the aperiodic offset and exponent parameters, the FOOOF algorithm also outputs parameters for periodic components by modelling individual oscillatory peaks of the 1/f-corrected power spectrum with a Gaussian (for details, see Donoghue et al., 2020). Periodic parameters include the center frequency of the peak, the peak’s height over and above the aperiodic component (i.e., aperiodic-adjusted peak power), and its bandwidth. In order to assess whether our procedure for deriving 1/f-corrected peak alpha characteristics impacted the outcomes of this study, comparisons between younger and older adults were repeated using FOOOF-derived center frequency and aperiodic-adjusted power for the largest peak in the 6–13 Hz extended alpha range (roughly equivalent to 1/f-corrected peak alpha frequency and power, respectively).

### 2.5 Statistics

Statistical analyses were performed using MATLAB (R2019a). Statistical significance was set at *p* < .05 (two-tailed), and data were checked for normality using the Kolmogorov-Smirnov test. Resting EEG power spectra (both uncorrected and 1/f-corrected) and the aperiodic component were averaged across all channels and compared between age groups using mixed-factorial ANOVA, with ‘age group’ (2 levels: younger and older) as between-subject factor and ‘frequency’ (77 levels: 2–40 Hz, in 0.5 Hz steps) as within-subject factor. *Post hoc* comparisons between age groups at each frequency step were performed using independent samples *t* tests, with the false-discovery rate (FDR) procedure used to correct for multiple comparisons (Curran-Everett, 2000). Age differences in aperiodic and peak alpha parameters were assessed in two ways. First, parameter values were averaged across all channels and compared between younger and older adults using Wilcoxon rank sum tests. Second, parameter values were compared between age groups at each channel using non-parametric cluster-based permutation statistics to control for multiple testing across channels (cluster threshold: *p* < .05, independent samples *t* test; test-statistic: maximum within-cluster summed *t* value; randomization: 5000 Monte-Carlo permutations) (Maris and Oostenveld, 2007). Cluster effect sizes were computed as the average Cohen’s *d* for all channels within the cluster. For both global average and cluster-based analyses, *p* values were Bonferroni-corrected for 6 comparisons (2 aperiodic parameters: offset and exponent; 4 peak alpha parameters: uncorrected peak alpha frequency, uncorrected peak alpha power, 1/f-corrected peak alpha frequency, and 1/f-corrected peak alpha power).

Lastly, the association between aperiodic activity, uncorrected peak alpha, and age were assessed by first using Spearman’s rank correlations between global average aperiodic and uncorrected peak alpha parameters in each age group separately (Bonferroni-corrected for 10 correlations). Correlation coefficients were transformed to z scores using Fisher’s transformation for comparisons between age groups. These were followed by binomial logistic regression analyses using aperiodic and uncorrected peak alpha parameters as predictor variables and age group (younger or older) as the outcome variable. This was included to further explore the confounding effect of aperiodic neural activity on the relationship between resting EEG peak alpha and age, examining the independent contributions of aperiodic and peak alpha parameters in predicting older age group membership. All *p* values reported in text and in figures have been Bonferroni-adjusted as required, unless otherwise indicated.

## 3. Results

### 3.1 Age differences in aperiodic offset and exponent

The aperiodic component averaged across all channels differed between age groups (age group × frequency: *F*_76,13300_ = 33.43, *p* < .001, η^2^_p_ = 0.16), with reduced power in the low-frequency range spanning delta, theta, alpha and low-beta frequencies (2–19 Hz) in older adults compared to younger adults (FDR-adjusted *p* ≤ .046) (Figure 2A). This age difference in aperiodic activity was reflected by a lower global average aperiodic offset (*Z* = −5.84, *p* = × 10^−8^, *r* = −0.44) (Figure 2B) and a smaller aperiodic exponent (i.e., flatter slope) (*Z* = − 5.34, *p* = 5.7 × 10^−7^, *r* = −0.40) (Figure 2C) in older adults. Cluster-based analyses similarly showed an age-related lowering of the aperiodic offset (max. summed *t* = −311.82, *p* = .002, *d* = −0.89) and exponent (max. summed *t* = −296.05, *p* = .002, *d* = −0.87), and suggest the age differences in aperiodic activity were not restricted to any particular region, with diffuse clusters extending across the whole scalp (Figures 2D–E).

**Figure 2.**
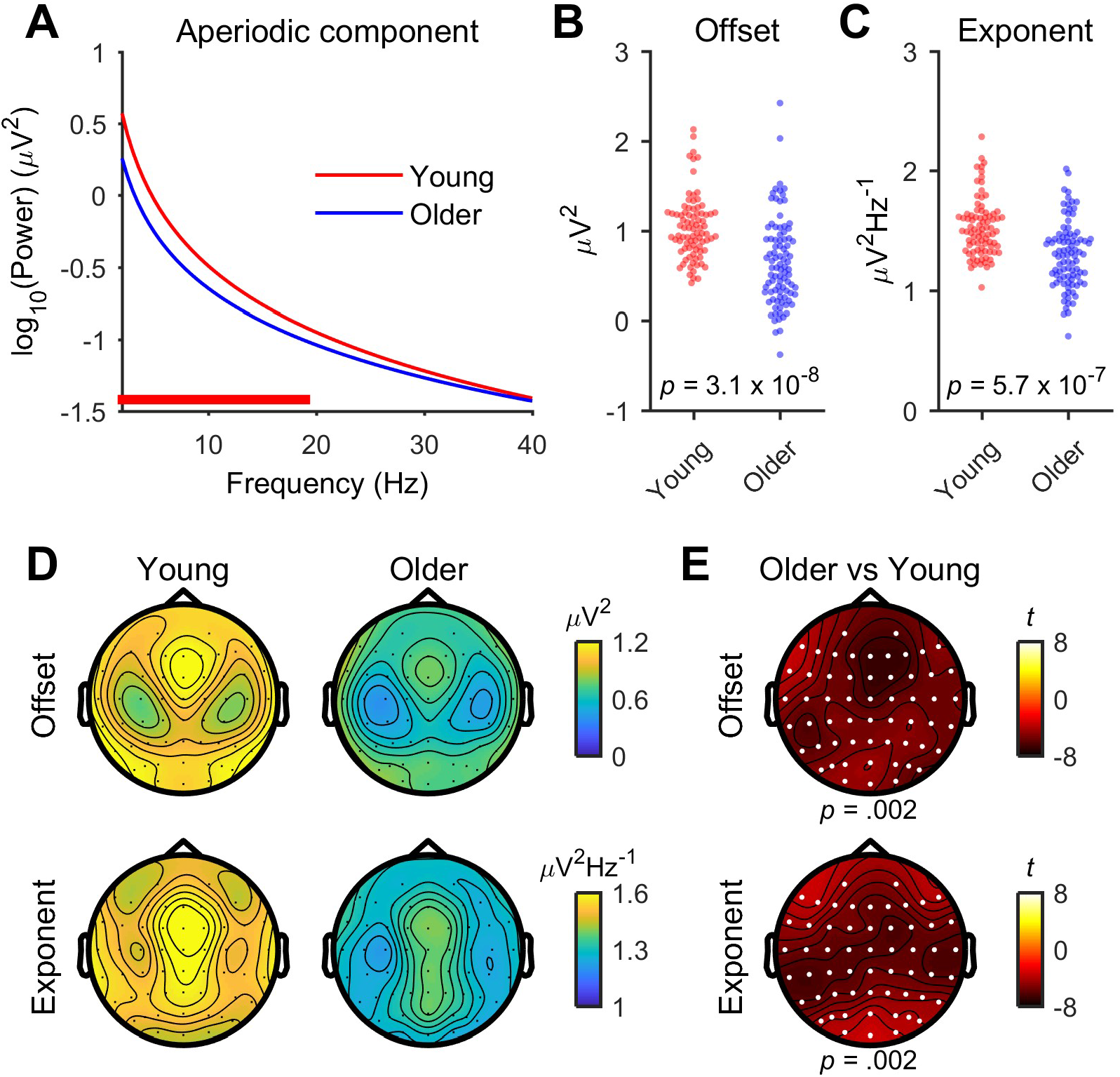
Age differences in resting EEG aperiodic neural activity. (A) The 1/f-like aperiodic component, as well as its (B) offset and (C) exponent parameters, averaged across all channels in younger and older adults. The thick red line above the x-axis in A indicates frequencies where logarithmic power differs between age groups (younger > older; FDR-adjusted *p* < .05; independent samples *t* test). (D) Topographical maps showing aperiodic offset and exponent for all channels in younger and older adults, and (E) *t*-statistic values comparing aperiodic offset and exponent between age groups. White dots in E indicate significant channels forming negative clusters (cluster-based permutation statistics).

### 3.2 Age differences in peak alpha before and after correcting for aperiodic activity

Figure 3 shows the influence of 1/f-like aperiodic neural activity on resting EEG power spectra and their peak alpha characteristics in younger and older adults, with results shown for uncorrected spectra on the left (Figures 3A_1_–E_1_) and 1/f-corrected spectra on the right (Figures 3A_2_–E_2_). Power spectra averaged across all channels differed between age groups, both for the uncorrected spectrum (age group × frequency: *F*_76,13300_ = 18.20, *p* < .001, η^2^_p_ = 0.094) (Figure 3A_1_) and after the aperiodic component was removed (age group × frequency: *F*_76,13300_ = 13.43, *p* < .001, η^2^_p_ = 0.071) (Figure 3A_2_). However, there were notable differences in the frequency bands affected and the direction of age effects. For the uncorrected spectrum, older adults had reduced power at delta, theta, and alpha frequencies (2–6.5 Hz and 9.5–12 Hz) (FDR-adjusted *p* ≤ .028) (Figure 3A_1_). Conversely, after correcting for aperiodic activity, they had reduced power at high-alpha frequencies (10–12 Hz) (FDR-adjusted *p* ≤ .043), but increased power at delta (3–3.5 Hz), low-alpha (8 Hz), beta (14–18.5 Hz and 22.5–28 Hz), and gamma frequencies (35.5 Hz) (FDR-adjusted *p* ≤ .043) (Figure 3A_2_).

**Figure 3.**
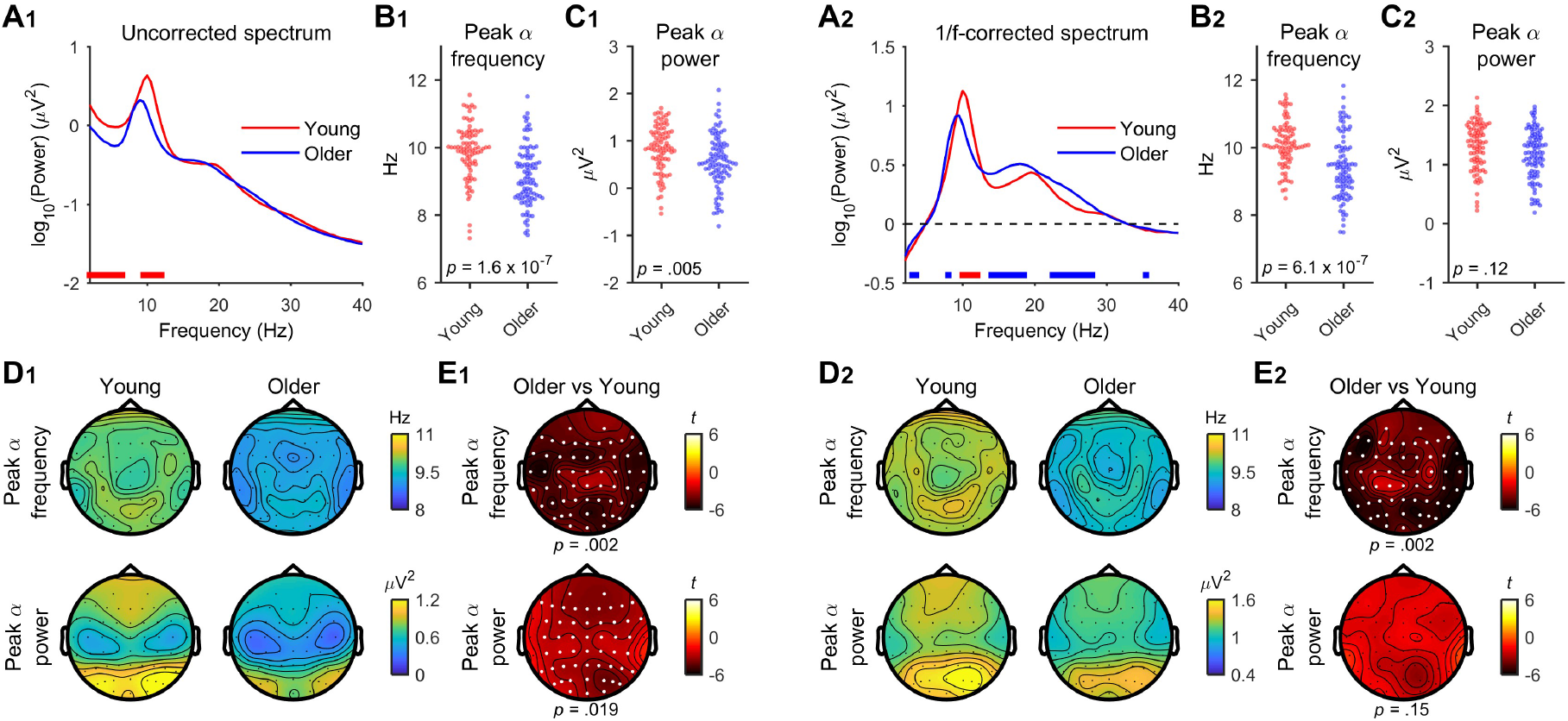
Age differences in resting EEG power spectra and their peak alpha characteristics. (A_1_, A_2_) The resting EEG power spectra, as well as their (B_1_, B_2_) peak alpha frequency and (C_1_, C_2_) peak alpha power, averaged across all channels in younger and older adults before (A_1_– C_1_) and after (A_2_–C_2_) correcting for the 1/f-like aperiodic component. The thick lines above the x-axis in A_1_ and A_2_ indicate frequencies where logarithmic power differs between age groups (red: younger > older, blue: older > younger; FDR-adjusted *p* < .05; independent samples t test). (D_1_, D_2_) Topographical maps showing uncorrected (D_1_) and 1/f-corrected (D_2_) peak alpha frequency and power for all channels in younger and older adults, and (E_1_, E_2_) *t*-statistic values comparing peak alpha frequency and power between age groups. White dots in E_1_/E_2_ indicate significant channels forming negative clusters (cluster-based permutation statistics).

Regarding peak alpha parameters averaged across all channels, older adults had slower global average peak alpha frequency than younger adults for both uncorrected (*Z* = −5.57, *p* = 1.6 × 10^−7^, *r* = −0.42) (Figure 3B_1_) and 1/f-corrected spectra (*Z* = −5.32, *p* = 6.1 × 10^−7^, *r* = − 0.40) (Figure 3B_2_). Although uncorrected peak alpha power was lower in older adults compared to younger adults (*Z* = −3.37, *p* = .005, *r* = −0.25) (Figure 3C_1_), this difference was no longer significant after removal of the aperiodic component (*Z* = −2.32, *p* = 0.12, *r* = −0.17) (Figure 3C_2_). The same can be seen using cluster-based analyses rather than averaging across all channels, with older adults having slower peak alpha frequency (max. summed *t* = −257.71, *p* = .002, *d* = −0.84) and lower peak alpha power (max. summed *t* = −164.14, *p* =.019, *d* = −0.51) before correcting for the aperiodic component (Figures 3D_1_–E_1_), as well as slower peak alpha frequency for the corrected spectrum (max. summed *t* = −260.32, *p* = .002, *d* = −0.84), but not differing in peak alpha power from younger adults after the aperiodic component was removed (max. summed *t* = −93.42, *p* = .15, *d* = −0.39) (Figures 3D_2_–E_2_).

Of note, none of these results were different when FOOOF-derived periodic parameters were used in place of 1/f-corrected peak alpha frequency and power, with slower alpha central frequency in older than younger adults, but no difference in aperiodic-adjusted peak alpha power between age groups (See Supplementary Figure 1).

### 3.3 Associations between aperiodic activity, uncorrected peak alpha, and age

To further explore the impact of aperiodic neural activity on resting EEG peak alpha in aging, we applied binomial logistic regressions to examine the independent contributions of aperiodic and peak alpha parameters in predicting age group membership. We first performed a series of Spearman’s rank correlations to characterize the associations between parameters in each age group separately (See Supplementary Figures 2 and 3). Averaged across all channels, aperiodic offset and exponent were strongly positively correlated in both younger (rho = 0.80, *p* = 4.5 × 10^−20^) and older adults (rho = 0.87, *p* = 1.3 × 10^−29^), with larger offset associated with a larger exponent (i.e., steeper slope) (Supplementary Figure 2). Aperiodic offset and uncorrected peak alpha frequency were moderately negatively correlated in younger adults (rho = −0.36, *p* = .007), with larger offset associated with slower peak alpha frequency (Supplementary Figure 3A). Although this relationship did not survive Bonferroni correction in older adults (rho = −0.26, *p* = .11), the correlation coefficients did not differ between age groups (*Z* = 0.70, *p* = .48). Offset and uncorrected peak alpha power were strongly positively correlated in both younger (rho = 0.50, *p* = 1.4 × 10^−5^) and older adults (rho = 0.53, *p* = 1.2 × 10^−6^) (Supplementary Figure 3B). Aperiodic exponent and uncorrected peak alpha frequency were moderately negatively correlated in both younger (rho = −0.42, *p* = 6.0 × 10^−4^) and older adults (rho = −0.32, *p* = .018), with a larger exponent associated with slower peak alpha frequency in both age groups (Supplementary Figure 3C). Larger exponents were also associated with higher peak alpha power in older (rho = 0.37, *p* = .003), but not younger adults (rho = 0.16, p = 1.00), although again, the correlation coefficients did not differ between age groups (*Z* = 1.47, *p* = .14) (Supplementary Figure 3D).

For binomial logistic regression, given the high collinearity between aperiodic offset and exponent, these were assessed as predictors in separate models (i.e., Models 2 and 3, respectively; see Table 1). First, we assessed uncorrected peak alpha frequency and power as predictor variables in a model without controlling for either aperiodic parameter (i.e., Model 1), and found that slower peak alpha frequency (*p* = 9.4 × 10^−7^) and reduced peak alpha power (*p* = .001) were both associated with older age (Table 1). When peak alpha frequency and power were both held constant, older age was associated with lower aperiodic offset (Model 2, *p* = 3.9 × 10^−7^) and smaller aperiodic exponent (Model 3, *p* = 1.6 × 10^−7^). Older age was also associated with slower peak alpha frequency after accounting for aperiodic offset (*p* = 3.7 × 10^−8^) and exponent (*p* = 1.1 × 10^−8^); however, peak alpha power was no longer associated with age group membership in either model (*p* ≥ .46). Thus, whereas altered aperiodic activity and slower peak alpha frequency were both strong independent predictors of older age, uncorrected peak alpha power was no longer associated with age after controlling for aperiodic offset and exponent.

**Table 1.** Results of binomial logistic regressions for predicting older age group membership.

| Model 1 | 95% CI |  |  |  |  |  |  |
| --- | --- | --- | --- | --- | --- | --- | --- |
|  | Estimate | SE | Z | p | Odds ratio | Lower | Upper |
| (Intercept) | 10.95 | 2.13 | 5.13 | $2.9 \times 10^{-7}$ | $5.7 \times 10^4$ | $8.6 \times 10^2$ | $3.7 \times 10^6$ |
| Peak $\alpha$ frequency | -1.06 | 0.22 | -4.90 | $9.4 \times 10^{-7}$ | 0.35 | 0.23 | 0.53 |
| Peak $\alpha$ power | -1.10 | 0.34 | -3.24 | .001 | 0.33 | 0.17 | 0.65 |
| Model 2 | 95% CI |  |  |  |  |  |  |
|  | Estimate | SE | Z | p | Odds ratio | Lower | Upper |
| (Intercept) | 16.46 | 2.78 | 5.92 | $3.3 \times 10^{-9}$ | $1.4 \times 10^7$ | $6.0 \times 10^4$ | $3.3 \times 10^9$ |
| Aperiodic offset | -3.23 | 0.64 | -5.07 | $3.9 \times 10^{-7}$ | 0.040 | 0.011 | 0.14 |
| Peak $\alpha$ frequency | -1.45 | 0.26 | -5.50 | $3.7 \times 10^{-8}$ | 0.23 | 0.14 | 0.39 |
| Peak $\alpha$ power | 0.34 | 0.45 | 0.74 | .46 | 1.40 | 0.58 | 3.40 |
| 95% CI |  |  |  |  |  |  |  |
| Model 3 | Estimate | SE | Z | p | Odds ratio | Lower | Upper |
| (Intercept) | 22.69 | 3.55 | 6.39 | $1.7 \times 10^{-10}$ | $7.2 \times 10^9$ | $6.8 \times 10^6$ | $7.6 \times 10^{12}$ |
| Aperiodic exponent | -5.23 | 1.00 | -5.24 | $1.6 \times 10^{-7}$ | 0.0054 | $7.6 \times 10^4$ | 0.038 |
| Peak $\alpha$ frequency | -1.57 | 0.28 | -5.71 | $1.1 \times 10^{-8}$ | 0.21 | 0.12 | 0.36 |
| Peak $\alpha$ power | -0.27 | 0.40 | -0.68 | .50 | 0.76 | 0.35 | 1.67 |

Note: CI = confidence interval; SE = standard error.

## 4. Discussion

In the present study, resting EEG recordings were used to investigate age differences in aperiodic activity, and establish if age differences in peak alpha characteristics were due to changes in the aperiodic component of the EEG power spectra. Consistent with previous findings, older adults had a reduced aperiodic offset and smaller aperiodic exponent than younger adults. Although peak alpha frequency and power were both reduced in older adults before correcting for aperiodic activity, only age differences in peak alpha frequency remained statistically significant after the aperiodic component was removed. This was supported by the results from binomial logistic regressions, which indicate slower peak alpha frequency, but not reduced peak alpha power, was able to independently predict older age group membership after controlling for the effects of aperiodic offset and exponent. Our findings provide further evidence that aperiodic neural activity changes with age, and should be controlled for in future studies measuring age-related changes in neural oscillatory activity.

### 4.1 The aperiodic component of EEG power spectra changes with age

Previous research has shown that the aperiodic component of the EEG power spectra changes with age. Voytek et al. (2015) examined the effect of age on aperiodic activity from two different datasets, including recordings from the cortical surface (electrocorticography; ECoG) in patients with intractable epilepsy while they performed an auditory task and scalp EEG recordings in healthy participants while they performed a visual working memory task. They observed that older age was associated with a flatter 1/f-like slope in both datasets and, moreover, the flattening of EEG power spectra with increased age statistically mediated age-related impairments in visual working memory performance. A similar age-related flattening of aperiodic activity has been observed for EEG recorded during language processing (Dave et al., 2018) and auditory discrimination tasks (Waschke et al., 2017), as well as for EEG recorded at rest (Donoghue et al., 2020; Kosciessa et al., 2020), with Donoghue et al. (2020) also reporting a lower aperiodic offset in older adults compared to younger adults. We also observed a reduction in both the aperiodic exponent and offset in older participants. These changes were observed across the scalp, and were not limited to any particular region. The results of the present study are thus in line with a growing body of literature demonstrating changes in aperiodic activity in older adults both at rest and during task performance.

There are several mechanisms which could underlie the observed age-related changes in aperiodic activity as measured with EEG. Evidence from empirical and computational studies has linked aperiodic features of electrophysiological power spectra to changes in population spiking statistics, with lower offsets and flatter slopes associated with slower firing rates and less synchronous spiking activity, respectively (Freeman and Zhai, 2009; Manning et al., 2009). According to the neural noise hypothesis, the reliability of neural communication diminishes with advanced age due to noisier (i.e., less synchronized) neural activity, contributing to age-related cognitive decline (Cremer and Zeef, 1987; Crossman and Szafran, 1956; Voytek and Knight, 2015). Thus, an age-related decrease in synchronous population spiking could explain the age differences in aperiodic activity observed here and in previous research.

Changes in the slope of electrophysiological power spectra may also be indicative of changes in synaptic excitation/inhibition balance. The co-regulation of excitatory and inhibitory output onto cortical neurons shapes the short-term dynamics within neuronal synapses, affecting the ability to form oscillations and efficient information transfer (Gao et al., 2017; Tatti et al., 2017). Changes in excitation/inhibition balance, over time, can have pathological effects in neuropsychiatric diseases such as autism, schizophrenia, as well as age-related neurodegenerative diseases, such as Alzheimer’s Disease (Cassani et al., 2018; Tatti et al., 2017; Voytek and Knight, 2015). Computational evidence supported by experimental data from rodents and non-human primates indicates that synaptic excitation/inhibition balance can be inferred from the aperiodic exponent of electrophysiological power spectra, with flatter slopes reflecting increased excitation and/or decreased inhibition (Gao et al., 2017). Although age-related increases in hippocampal excitation/inhibition ratio have been suggested from some studies using animal models (El-Hayek et al., 2013; Wilson et al., 2005), other studies in humans have reported no change (Noda et al., 2017) or lower excitation/inhibition ratios in older adults (Legon et al., 2016). Thus, the possible contribution of altered excitation/inhibition balance to age differences in aperiodic activity remains speculative and requires further research.

Maintaining functional neural activity throughout the lifespan is an integral part of healthy aging, therefore, it is essential to investigate other features of brain physiology that could explain differences in aperiodic activity between younger and older adults. For example, Muthukumaraswamy and Liley (2018) showed that the 1/f-like nature of electrophysiological power spectra could be modelled using a collection of damped oscillators with a distribution of relaxation rates. Changes in the properties of the neural generators proposed to underlie these damped oscillators could underlie age-related changes in aperiodic activity. Conversely, age differences could emerge from between-group differences in the physical properties of brain tissue that need not necessarily be due to neural processes (e.g., Bedard et al., 2006; Bedard and Destexhe, 2009). Consequently, additional research into the neural and non-neural factors that drive 1/f scaling of EEG power spectra, and how these factors change with age, is required.

### 4.2 Advanced age decreases peak alpha frequency, but not power

In support of previous findings (Bablioni et al., 2006; Grandy et al., 2013; Klimesch, 1999; Scally et al., 2018; Sghirripa et al., 2020), this study provides more evidence that older adults have slower peak alpha frequency compared to younger adults, even after accounting for age differences in aperiodic activity. Age-related peak alpha slowing may be due to various neuroanatomical changes in the aging brain. Thalamo-cortical circuits have been associated with alpha oscillatory activity (Hughes and Crunelli, 2005; Olejniczak, 2006; Schaul, 1998), and biophysical models have shown peak alpha frequency to be sensitive to changes in conduction delays in thalamo-cortical feedback loops (Roberts and Robinson, 2008; Robinson et al., 2001). Accordingly, empirical data has shown that individual differences in peak alpha frequency are related to white matter structure (Valdés-Hernández et al., 2010), pointing to white matter deterioration in thalamo-cortical circuits as a potential factor explaining age-related slowing of alpha oscillations.

Unlike our results for peak alpha frequency, age differences in peak alpha power were not statistically significant after accounting for the aperiodic component. The confounding effect of aperiodic activity on alpha oscillatory power has been observed previously in a recent study by Ouyang et al. (2020), which used structural equation modelling to investigate the relationships between aperiodic activity, alpha power, and cognitive processing speed in healthy young adults. While an initial model was suggestive of an association between eyes-open alpha power and cognitive processing speed, this relationship was no longer present in subsequent models dissociating alpha oscillations from aperiodic activity. Regarding alpha power and aging, Donoghue et al. (2020) assessed age differences in resting EEG alpha after isolating the oscillatory peak from the aperiodic background. Although the age difference in aperiodic-adjusted peak alpha power was reduced compared to non-adjusted power, a statistically significant difference between younger and older adults was still present. Our findings from a larger sample using whole-scalp EEG and multiple analysis approaches could not find evidence for an age effect on peak alpha power after accounting for aperiodic activity. These results further highlight the importance of accounting for aperiodic activity when analyzing age-related spectral power changes.

### 4.3 Conclusions

The present study has demonstrated age differences in aperiodic activity measured using resting EEG in a large sample of healthy younger and older adults. Compared to young adults, older adults had reduced aperiodic offset and smaller aperiodic exponent. After correcting for aperiodic activity, age differences in peak alpha frequency persisted, while peak alpha power was no longer statistically different between age groups. Together, our findings provide further evidence that the aperiodic component of the resting EEG signal is altered with aging and should be considered when investigating neural oscillatory activity.

## Supplementary material

**Supplementary Figure 1.**
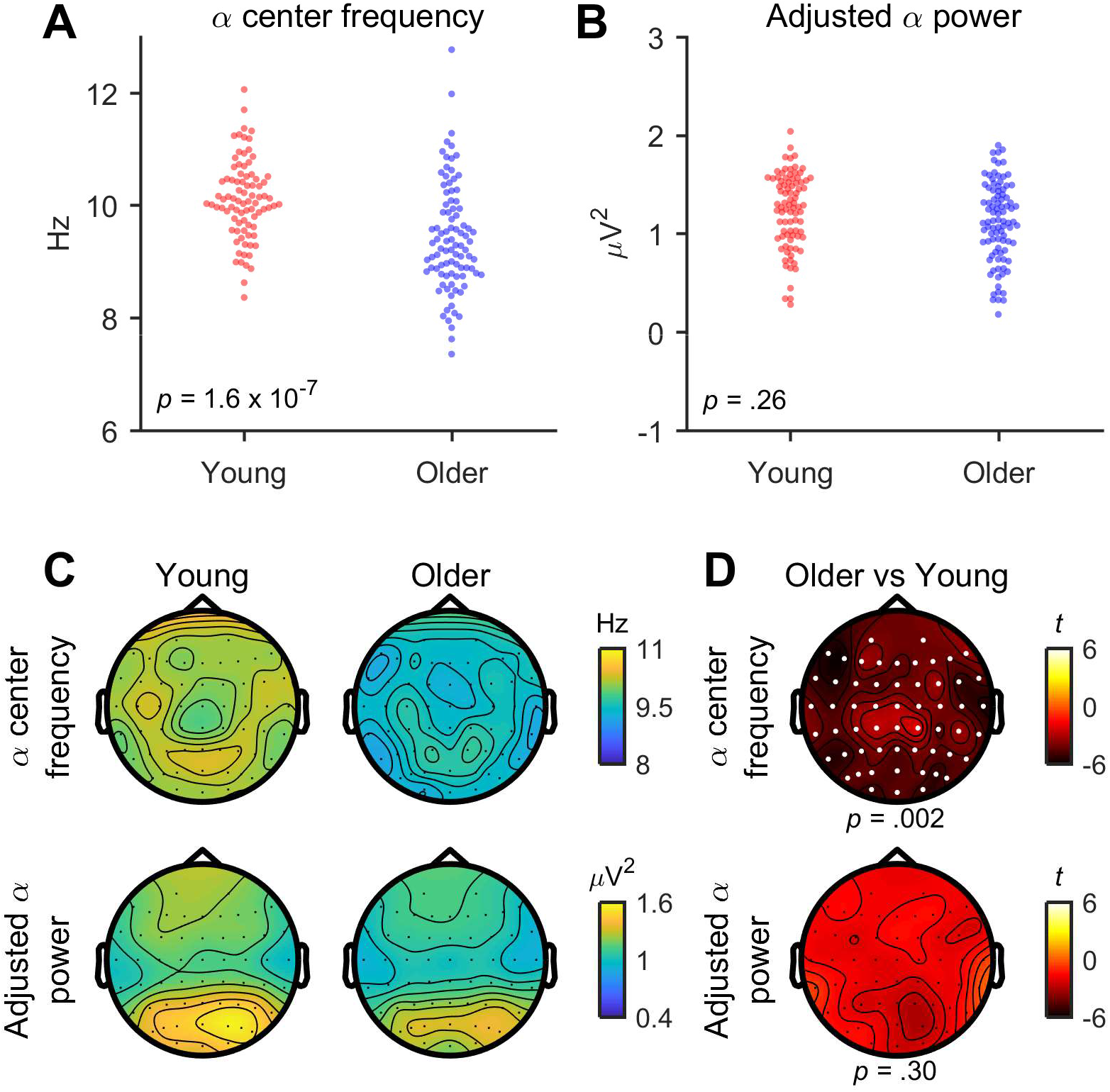
Age differences in FOOOF-derived periodic alpha parameters. When averaged across all channels, older adults had (A) slower alpha center frequency than younger adults (*Z* = −5.37, *p* = 1.6 × 10^−7^, *r* = −0.40); however, (B) aperiodic-adjusted alpha power did not differ between age groups (*Z* = −2.02, *p* = .26, *r* = −0.15). (C) Topographical maps showing alpha center frequency and aperiodic-adjusted alpha power for all channels in younger and older adults, and (D) *t*-statistic values comparing alpha center frequency and aperiodic-adjusted alpha power between age groups. White dots in D indicate significant channels forming negative clusters (cluster-based permutation statistics).

**Supplementary Figure 2.**
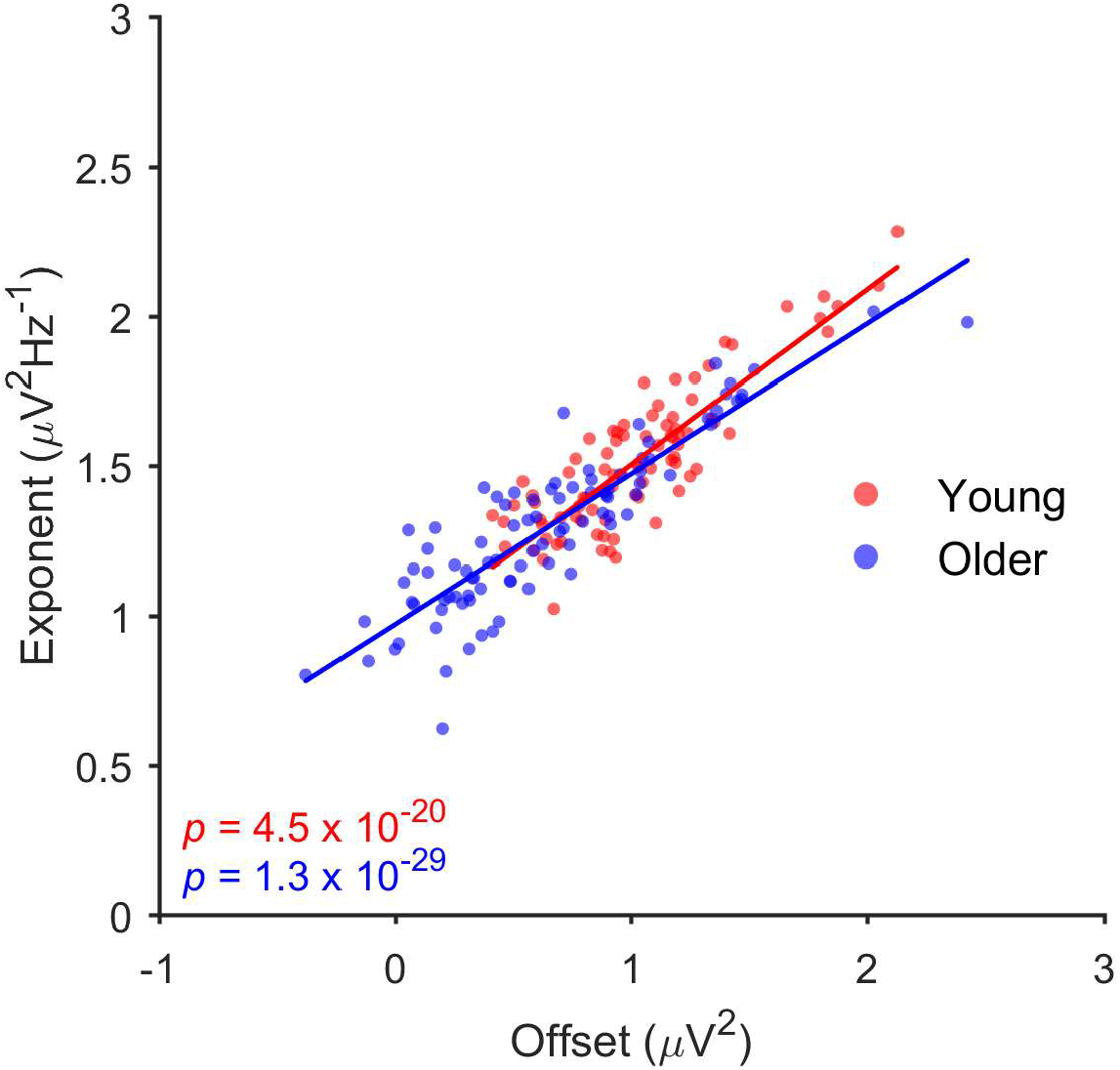
Spearman’s rank correlations between aperiodic offset and exponent, averaged across all channels, in younger and older adults.

**Supplementary Figure 3.**
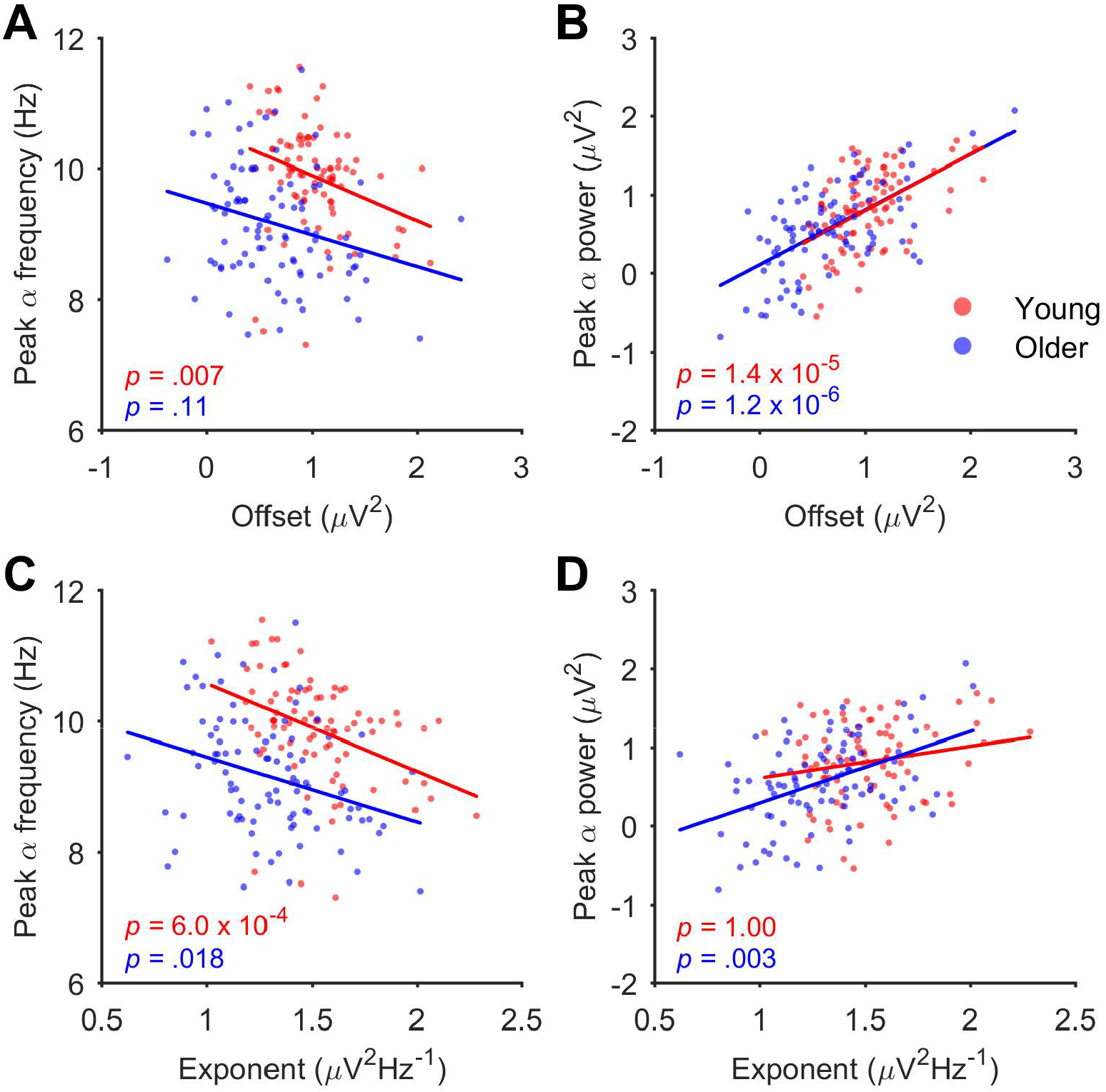
Spearman’s rank correlations between aperiodic parameters (A, B: offset; C, D: exponent) and uncorrected peak alpha parameters (A, C: peak alpha frequency; B, D: peak alpha power), averaged across all channels, in younger and older adults.

## References

Babiloni, C., Binetti, G., Cassarino, A., Dal Forno, G., Del Percio, C., Ferreri, F., Ferri, R., Frisoni, G., Galderisi, S., Hirata, K., Lanuzza, B., Miniussi, C., Mucci, A., Nobili, F., Rodriguez, G., Romani, G.L., Rossini, P.M., 2006. Sources of cortical rhythms in adults during physiological aging: A multicentric EEG study. Hum. Brain Mapp. 27, 162–172. https://doi.org/10.1002/hbm.20175

Bédard, C., Destexhe, A., 2009. Macroscopic Models of Local Field Potentials and the Apparent 1/f Noise in Brain Activity. Biophysical Journal 96, 2589–2603. https://doi.org/10.1016/j.bpj.2008.12.3951

Bédard, C., Kröger, H., Destexhe, A., 2006. Does the 1/f Frequency Scaling of Brain Signals Reflect Self-Organized Critical States? Phys. Rev. Lett. 97, 118102. https://doi.org/10.1103/PhysRevLett.97.118102

Cassani, R., Estarellas, M., San-Martin, R., Fraga, F.J., Falk, T.H., 2018. Systematic Review on Resting-State EEG for Alzheimer’s Disease Diagnosis and Progression Assessment. Dis Markers 2018, 5174815. https://doi.org/10.1155/2018/5174815

Colloby, S.J., Cromarty, R.A., Peraza, L.R., Johnsen, K., Jóhannesson, G., Bonanni, L., Onofrj, M., Barber, R., O’Brien, J.T., Taylor, J.-P., 2016. Multimodal EEG-MRI in the differential diagnosis of Alzheimer’s disease and dementia with Lewy bodies. J. Psychiatr. Res. 78, 48–55. https://doi.org/10.1016/j.jpsychires.2016.03.010

Cremer, R., Zeef, E.J., 1987. What Kind of Noise Increases With Age? J. Gerontol. 42, 515–518. https://doi.org/10.1093/geronj/42.5.515

Crossman, E.R., Szafran, J., 1956. Changes with age in the speed of information-intake and discrimination. Experientia 128–34; discussion, 135.

Curran-Everett, D., 2000. Multiple comparisons: philosophies and illustrations. Am. J. Physiol. Regul. Integr. Comp. Physiol. 279, R1–R8. https://doi.org/10.1152/ajpregu.2000.279.1.R1

Dave, S., Brothers, T.A., Swaab, T.Y., 2018. 1/f neural noise and electrophysiological indices of contextual prediction in aging. Brain Res. 1691, 34–43. https://doi.org/10.1016/j.brainres.2018.04.007

Delorme, A., Makeig, S., 2004. EEGLAB: an open source toolbox for analysis of single-trial EEG dynamics including independent component analysis. J. Neurosci. Methods 134, 9–21. https://doi.org/10.1016/j.jneumeth.2003.10.009

Donoghue, T., Haller, M., Peterson, E.J., Varma, P., Sebastian, P., Gao, R., Noto, T., Lara, A.H., Wallis, J.D., Knight, R.T., Shestyuk, A., Voytek, B., 2020. Parameterizing neural power spectra into periodic and aperiodic components. Nat. Neurosci. 23, 1655–1665. https://doi.org/10.1038/s41593-020-00744-x

El-Hayek, Y.H., Wu, C., Ye, H., Wang, J., Carlen, P.L., Zhang, L., 2013. Hippocampal excitability is increased in aged mice. Exp. Neurol. 247, 710–719. https://doi.org/10.1016/j.expneurol.2013.03.012

Folstein, M.F., Folstein, S.E., McHugh, P.R., 1975. “Mini-mental state”: A practical method for grading the cognitive state of patients for the clinician. J. Psychiatr. Res. 12, 189–198. https://doi.org/10.1016/0022-3956(75)90026-6

Freeman, W.J., Zhai, J., 2009. Simulated power spectral density (PSD) of background electrocorticogram (ECoG). Cogn. Neurodyn. 3, 97–103. https://doi.org/10.1007/s11571-008-9064-y

Gao, R., Peterson, E.J., Voytek, B., 2017. Inferring synaptic excitation/inhibition balance from field potentials. Neuroimage 158, 70–78. https://doi.org/10.1016/j.neuroimage.2017.06.078

Garn, H., Coronel, C., Waser, M., Caravias, G., Ransmayr, G., 2017. Differential diagnosis between patients with probable Alzheimer’s disease, Parkinson’s disease dementia, or dementia with Lewy bodies and frontotemporal dementia, behavioral variant, using quantitative electroencephalographic features. J. Neural Transm. 124, 569–581. https://doi.org/10.1007/s00702-017-1699-6

Grandy, T.H., Werkle-Bergner, M., Chicherio, C., Schmiedek, F., Lövdén, M., Lindenberger, U., 2013. Peak individual alpha frequency qualifies as a stable neurophysiological trait marker in healthy younger and older adults. Psychophysiology 50, 570–582. https://doi.org/10.1111/psyp.12043

Grigolini, P., Aquino, G., Bologna, M., Luković, M., West, B.J., 2009. A theory of 1/f noise in human cognition. Phys. A: Stat. Mech. Appl. 388, 4192–4204. https://doi.org/10.1016/j.physa.2009.06.024

Harada, C.N., Natelson Love, M.C., Triebel, K., 2013. Normal Cognitive Aging. Clin. Geriatr. Med. 29, 737–752. https://doi.org/10.1016/j.cger.2013.07.002

Hughes, S.W., Crunelli, V., 2005. Thalamic Mechanisms of EEG Alpha Rhythms and Their Pathological Implications. Neuroscientist 11, 357–372. https://doi.org/10.1177/1073858405277450

Hyvärinen, A., Oja, E., 2000. Independent component analysis: algorithms and applications. Neural. Netw. 13, 411–430. https://doi.org/10.1016/s0893-6080(00)00026-5

Jeong, D.-H., Kim, Y.-D., Song, I.-U., Chung, Y.-A., Jeong, J., 2016. Wavelet Energy and Wavelet Coherence as EEG Biomarkers for the Diagnosis of Parkinson’s Disease-Related Dementia and Alzheimer’s Disease. Entropy 18, 8. https://doi.org/10.3390/e18010008

Klimesch, W., 1999. EEG alpha and theta oscillations reflect cognitive and memory performance: a review and analysis. Brain Res. Rev. 29, 169–195. https://doi.org/10.1016/S0165-0173(98)00056-3

Kosciessa, J.Q., Kloosterman, N.A., Garrett, D.D., 2020. Standard multiscale entropy reflects neural dynamics at mismatched temporal scales: What’s signal irregularity got to do with it? PLoS Comput. Biol. 16, e1007885. https://doi.org/10.1371/journal.pcbi.1007885

Kumar, P., Parmananda, P., 2018. Control, synchronization, and enhanced reliability of aperiodic oscillations in the Mercury Beating Heart system. Chaos 28, 045105. https://doi.org/10.1063/1.5006697

Legon, W., Punzell, S., Dowlati, E., Adams, S.E., Stiles, A.B., Moran, R.J., 2016. Altered Prefrontal Excitation/Inhibition Balance and Prefrontal Output: Markers of Aging in Human Memory Networks. Cereb. Cortex 26, 4315–4326. https://doi.org/10.1093/cercor/bhv200

Mahjoory, K., Schoffelen, J.-M., Keitel, A., Gross, J., 2020. The frequency gradient of human resting-state brain oscillations follows cortical hierarchies. eLife 9, e53715. https://doi.org/10.7554/eLife.53715

Manning, J.R., Jacobs, J., Fried, I., Kahana, M.J., 2009. Broadband Shifts in Local Field Potential Power Spectra Are Correlated with Single-Neuron Spiking in Humans. J. Neurosci. 29, 13613–13620. https://doi.org/10.1523/JNEUROSCI.2041-09.2009

Maris, E., Oostenveld, R., 2007. Nonparametric statistical testing of EEG- and MEG-data. J. Neurosci. Methods 164, 177–190. https://doi.org/10.1016/j.jneumeth.2007.03.024

Michels, L., Muthuraman, M., Lüchinger, R., Martin, E., Anwar, A.R., Raethjen, J., Brandeis, D., Siniatchkin, M., 2013. Developmental changes of functional and directed resting-state connectivities associated with neuronal oscillations in EEG. NeuroImage 81, 231–242. https://doi.org/10.1016/j.neuroimage.2013.04.030

Miller, K.J., Hermes, D., Honey, C.J., Hebb, A.O., Ramsey, N.F., Knight, R.T., Ojemann, J.G., Fetz, E.E., 2012. Human Motor Cortical Activity Is Selectively Phase-Entrained on Underlying Rhythms. PLoS Comput. Biol. 8, e1002655. https://doi.org/10.1371/journal.pcbi.1002655

Mioshi, E., Dawson, K., Mitchell, J., Arnold, R., Hodges, J.R., 2006. The Addenbrooke’s Cognitive Examination Revised (ACE-R): a brief cognitive test battery for dementia screening. Int. J. Geriatr. Psychiatry 21, 1078–1085. https://doi.org/10.1002/gps.1610

Muthukumaraswamy, S.D., Liley, D.TJ., 2018. 1/f electrophysiological spectra in resting and drug-induced states can be explained by the dynamics of multiple oscillatory relaxation processes. NeuroImage 179, 582–595. https://doi.org/10.1016/j.neuroimage.2018.06.068

Neto, E., Allen, E.A., Aurlien, H., Nordby, H., Eichele, T., 2015. EEG Spectral Features Discriminate between Alzheimer’s and Vascular Dementia. Front. Neurol. 6, 25. https://doi.org/10.3389/fneur.2015.00025

Neto, E., Biessmann, F., Aurlien, H., Nordby, H., Eichele, T., 2016. Regularized Linear Discriminant Analysis of EEG Features in Dementia Patients. Front. Aging Neurosci. 8, 273. https://doi.org/10.3389/fnagi.2016.00273

Nishida, K., Yoshimura, M., Isotani, T., Yoshida, T., Kitaura, Y., Saito, A., Mii, H., Kato, M., Takekita, Y., Suwa, A., Morita, S., Kinoshita, T., 2011. Differences in quantitative EEG between frontotemporal dementia and Alzheimer’s disease as revealed by LORETA. Clin. Neurophysiol. 122, 1718–1725. https://doi.org/10.1016/j.clinph.2011.02.011

Noda, Y., Zomorrodi, R., Cash, R.F.H., Barr, M.S., Farzan, F., Rajji, T.K., Chen, R., Daskalakis, Z.J., Blumberger, D.M., 2017. Characterization of the influence of age on GABAA and glutamatergic mediated functions in the dorsolateral prefrontal cortex using paired-pulse TMS-EEG. Aging 9, 556–567. https://doi.org/10.18632/aging.101178

Olejniczak, P., 2006. Neurophysiologic Basis of EEG. J. Clin. Neurophysiol. 23, 186–189. https://doi.org/10.1097/01.wnp.0000220079.61973.6c

Oostenveld, R., Fries, P., Maris, E., Schoffelen, J.-M., 2011. FieldTrip: Open Source Software for Advanced Analysis of MEG, EEG, and Invasive Electrophysiological Data. Comput. Intell. Neurosci. 2011, 156869. https://doi.org/10.1155/2011/156869

Ouyang, G., Hildebrandt, A., Schmitz, F., Herrmann, C.S., 2020. Decomposing alpha and 1/f brain activities reveals their differential associations with cognitive processing speed. NeuroImage 205, 116304. https://doi.org/10.1016/j.neuroimage.2019.116304

Phillips, L.H., Andrés, P., 2010. The cognitive neuroscience of aging: New findings on compensation and connectivity. Cortex, The Cognitive Neuroscience of Aging 46, 421–424. https://doi.org/10.1016/j.cortex.2010.01.005

Roberts, J.A., Robinson, P.A., 2008. Modeling distributed axonal delays in mean-field brain dynamics. Phys. Rev. E 78, 051901. https://doi.org/10.1103/PhysRevE.78.051901

Robinson, P.A., Loxley, P.N., O’Connor, S.C., Rennie, C.J., 2001. Modal analysis of corticothalamic dynamics, electroencephalographic spectra, and evoked potentials. Phys. Rev. E 63, 041909. https://doi.org/10.1103/PhysRevE.63.041909

Rogasch, N.C., Sullivan, C., Thomson, R.H., Rose, N.S., Bailey, N.W., Fitzgerald, P.B., Farzan, F., Hernandez-Pavon, J.C., 2017. Analysing concurrent transcranial magnetic stimulation and electroencephalographic data: A review and introduction to the open-source TESA software. NeuroImage 147, 934–951. https://doi.org/10.1016/j.neuroimage.2016.10.031

Scally, B., Burke, M.R., Bunce, D., Delvenne, J.-F., 2018. Resting-state EEG power and connectivity are associated with alpha peak frequency slowing in healthy aging. Neurobiol. Aging 71, 149–155. https://doi.org/10.1016/j.neurobiolaging.2018.07.004

Schaul, N., 1998. The fundamental neural mechanisms of electroencephalography. Electroencephalogr. Clin. Neurophysiol. 106, 101–107. https://doi.org/10.1016/S0013-4694(97)00111-9

Schaworonkow, N., Voytek, B., 2021. Longitudinal changes in aperiodic and periodic activity in electrophysiological recordings in the first seven months of life. Dev. Cogn. Neurosci. 47, 100895. https://doi.org/10.1016/j.dcn.2020.100895

Sghirripa, S., Graetz, L., Merkin, A., Rogasch, N.C., Semmler, J.G., Goldsworthy, M.R., 2021. Load-dependent modulation of alpha oscillations during working memory encoding and retention in young and older adults. Psychophysiology 58, e13719. https://doi.org/10.1111/psyp.13719

Tatti, R., Haley, M.S., Swanson, O.K., Tselha, T., Maffei, A., 2017. Neurophysiology and Regulation of the Balance Between Excitation and Inhibition in Neocortical Circuits. Biol. Psychiatry, Cortical Excitation-Inhibition Balance and Dysfunction in Psychiatric Disorders 81, 821–831. https://doi.org/10.1016/j.biopsych.2016.09.017

Vaden, R.J., Hutcheson, N.L., McCollum, L.A., Kentros, J., Visscher, K.M., 2012. Older adults, unlike younger adults, do not modulate alpha power to suppress irrelevant information. NeuroImage 63, 1127–1133. https://doi.org/10.1016/j.neuroimage.2012.07.050

Valdés-Hernández, P.A., Ojeda-González, A., Martínez-Montes, E., Lage-Castellanos, A., Virués-Alba, T., Valdés-Urrutia, L., Valdes-Sosa, P.A., 2010. White matter architecture rather than cortical surface area correlates with the EEG alpha rhythm. NeuroImage 49, 2328–2339. https://doi.org/10.1016/j.neuroimage.2009.10.030

Voytek, B., Knight, R.T., 2015. Dynamic Network Communication as a Unifying Neural Basis for Cognition, Development, Aging, and Disease. Biol. Psychiatry, Cortical Oscillations for Cognitive/Circuit Dysfunction in Psychiatric Disorders 77, 1089–1097. https://doi.org/10.1016/j.biopsych.2015.04.016

Voytek, B., Kramer, M.A., Case, J., Lepage, K.Q., Tempesta, Z.R., Knight, R.T., Gazzaley, A., 2015. Age-Related Changes in 1/f Neural Electrophysiological Noise. J. Neurosci. 35, 13257–13265. https://doi.org/10.1523/JNEUROSCI.2332-14.2015

Waschke, L., Wöstmann, M., Obleser, J., 2017. States and traits of neural irregularity in the age-varying human brain. Sci. Rep. 7, 17381. https://doi.org/10.1038/s41598-017-17766-4

Wilson, I.A., Ikonen, S., Gallagher, M., Eichenbaum, H., Tanila, H., 2005. Age-Associated Alterations of Hippocampal Place Cells Are Subregion Specific. J. Neurosci. 25, 6877–6886. https://doi.org/10.1523/JNEUROSCI.1744-05.2005

Winawer, J., Kay, K.N., Foster, B.L., Rauschecker, A.M., Parvizi, J., Wandell, B.A., 2013. Asynchronous Broadband Signals Are the Principal Source of the BOLD Response in Human Visual Cortex. Curr. Biol. 23, 1145–1153. https://doi.org/10.1016/j.cub.2013.05.001

